# Polarization affects viscoelasticity and podosome distribution of primary human macrophages

**DOI:** 10.64898/2026.10.26.759594

**Authors:** Hasnae El Showk, Chiara Lunardi, Riduan Erkan, Nicholas A. Kurniawan, Anthal I.P.M Smits, Livia Angeloni

## Abstract

Macrophages are essential cells of the innate immune system that regulate wound healing and tissue homeostasis. They exert diverse functions, depending on their polarization state, which can lead to either functional healing of the tissue or fibrosis. Macrophage polarization state is affected by both the local biochemical and mechanical microenvironment within the tissue, suggesting a possible dependence of macrophage function on cell mechanical state, which has been poorly explored to date. In this study, we provide a systematic and quantitative characterization of the viscoelastic properties of unstimulated, pro-inflammatory, and anti-inflammatory monocyte-derived primary human macrophages at the single-cell level using atomic force microscopy. Pro-inflammatory macrophages exhibited a lower elastic modulus (E = 458.8 ± 179.9 Pa) and a more dissipative response (low frequency exponent *α* = 0.22 ± 0.03) than anti-inflammatory macrophages (E = 716 ± 345.6 Pa and *α* = 0.15 ± 0.04). These differences in mechanical properties were independent of cell morphology but correlated with differences in podosome distribution with pro-inflammatory macrophages exhibiting a dispersed distribution, and anti-inflammatory macrophages a more clustered distribution. This study identifies polarization-dependent mechanical differences in human primary macrophages and provides a quantitative experimental framework for future work in macrophage mechanobiology.

## 1 Introduction

Macrophages are cells of the innate immune system that play a pivotal role in modulating inflammation and tissue repair. They are characterized by their ability to polarize into a spectrum of phenotypes ranging from a pro-inflammatory phenotype (termed M1) to an anti-inflammatory phenotype (termed M2). They are known to be highly plastic and sensitive cells that dynamically alter their phenotype in response to biochemical, mechanical, and physical signals in their surrounding environment^1^. While the interaction between immune cells and their environment has been extensively studied, the mechanical properties of these cells are often overlooked. In the last few years, the study of cell mechanics has attracted growing interest as evidence links it closely to cell function and to the understanding of disease progression ^2,3^. For instance, it is widely established that cancer cells have, on average, a lower elastic modulus compared to healthy cells from the same tissue. Similarly, a decrease in the loss tangent (the phase angle between stress and strain) has been associated with an increase in metastatic potential, suggesting that increased fluidity of the cytoskeleton may enhance cell invasiveness ^4^. Recently, emerging evidence on macrophages’ mechanosensitivity has raised the hypothesis of a direct relationship between their biological behavior and mechanical state. For instance, macrophage activation in response to soluble factors and physical cues correlates with actin polymerization ^5,6^ and membrane tension ^7^, which have been shown to be primary contributors to cell mechanical properties for several mammalian cell types ^8^. Furthermore, it has also been suggested that actin cytoskeletal remodeling generates contractile forces that contribute to the functional activation of macrophages through the translocation of transcription factors to the nucleus, such as MRTF-A, YAP, and NF-*κβ*, which can facilitate changes in gene transcriptions correlated with macrophage function ^9^. In recent work, Ni et al. demonstrated a correlation between adhesion, membrane deformability and macrophage activation in response to substrate stiffness. Specifically, the authors hypothesized that macrophages are activated in response to changes in their mechanical environment via an adhesion-mediated plasma membrane deformation mechanism, which might depend on cell mechanical properties (e.g. membrane tension) and, in turn, regulates cytoskeletal remodeling ^7^. This data support the hypothesis that macrophage mechanical properties are correlated with their function. However, the specific causal relationship among macrophage mechanical properties, cytoskeletal organization, and function remains an open fundamental question in macrophage mechanobiology.

Understanding the relationship between macrophage mechanics and function could improve our ability to predict how macrophages will respond to complex environments (tissue or biomaterial) and pave the way for the development of advanced therapeutic strategies, such as targeting cytoskeletal mechanics to reprogram macrophage behavior. However, the mechanical properties of primary human macrophages have been poorly characterized to date. This lack of data can be attributed to the high phenotypic plasticity of macrophages which makes them more sensitive than other cell types to specific culture conditions, and therefore difficult to integrate into live-cell mechanical testing approaches. As a consequence, there is a need for reliable methods capable of quantitatively characterizing the mechanical properties of living macrophages under biologically relevant conditions. Few studies have conducted mechanical characterization of mouse- or cell-line-derived macrophages ^10,11^. However, mouse models and macrophage-like cell lines do not accurately reflect human immune biology. To the best of our knowledge, only four studies in literature have focused on the mechanical characterization of monocyte-derived primary human macrophages ^12–15^. Evers et al. applied optical tweezers (OT) and acoustic force spectroscopy (AFS) to study the viscoelastic properties of monocyte-derived primary human macrophages, concluding that anti-inflammatory macrophages appear to exhibit a higher elastic modulus (600 vs 800 Pa) and a more “solid-like” behavior, which was associated with a farther, straighter, and faster migration than pro-inflammatory macrophages ^13^. Although efficient for high-throughput mechanical characterization of suspended or weakly adhered cells, OT and AFS preclude quantification of the viscoelastic properties of macrophages adhering on substrates of interest. Atomic force microscopy (AFM)-based techniques enable quantitative mechanical analysis of adherent cells and were used to measure the elastic modulus of primary human monocyte-derived macrophages under different conditions. As an example, Leporatti and colleagues used colloidal force microscopy to measure the elastic modulus of unstimulated and LPS-stimulated primary human macrophages and observed a softening of the cell upon LPS stimulation with Young modulus values of 290 and 830 Pa for LPS-stimulated and unstimulated macrophages, respectively ^14^. On the contrary, Chen and colleagues measured the elastic modulus of unstimulated, pro- and anti-inflammatory primary macrophages using AFM Force Mapping (spherical probe, r=10 *µ*m), finding elastic modulus values significantly lower in comparison to the above-mentioned studies, with an average of 165 to ≈ 225 Pa, with the pro-inflammatory phenotype slightly stiffer (with no statistical difference) than unstimulated and anti-inflammatory macrophages ^15^. Crucially, these AFM studies did not include the viscoelastic nature of cells, focusing only on the quantification of the elastic modulus and therefore providing only a partial description of the mechanical behavior of macrophages. Nevertheless, despite substantial differences in experimental conditions, stimulation factors, and mechanical measurement approaches, these studies consistently suggest that pro- and anti-inflammatory macrophages exhibit distinct mechanical properties, potentially reflecting differences in their biological and functional roles. In this work, we used AFM to quantify the elastic modulus and viscoelastic behavior of adherent primary human monocyte-derived macrophages, combining force spectroscopy and microrheology. We applied this procedure to biochemically stimulated pro-inflammatory and anti-inflammatory phenotypes to investigate differences in cell mechanical properties that may correlate with macrophage function.

## 2 Materials and Methods

### 2.1 Human monocyte isolation

Buffy coats from healthy anonymized donors(Supplementary Table.**??**)were purchased from Sanquin with informed consent and approval from the Institutional Medical Ethical Committee of Sanquin Research. Human peripheral blood mononucleated cells (hPBMCs) were isolated by density gradient centrifugation with Lymphoprep (StemCell) and monocytes were isolated using the plastic adhesion method. Briefly, 100 million hPBMCs were seeded in a T-75 suspension cell culture flask (Greiner-Bio) and incubated for 1h (5% CO_2_, 37°C) in 10 ml of RPMI medium supplemented with 1% penicillin/streptomycin (Lonza), 2% Human Serum (heat-inactivated, pooled AB male donors; Sanquin), and 20 ng/ml Macrophage colony-stimulating factor (M-CSF; Peprotech) to allow for monocyte adhesion. The medium was then refreshed, discarding any non-adherent cells.

### 2.2 Macrophage polarization and culture

Isolated monocytes were cultured in the presence of M-CSF for 6 days to differentiate into macrophages (MΦ). Cells were then detached by incubating (5% CO_2_, 37°C) with Accutase (StemCell) for 45 minutes and seeded for either RT-qPCR, immunofluorescence (IF) imaging or AFM at a density of 35,000 cells per cm^2^. Cells were cultured in a 24-well plate for qPCR, a 24-well plate with a 13mm glass coverslip for IF imaging, and in a glass-bottom 35mm Petri dish (Mattek) for AFM measurements. Macrophages were incubated for 24h in the presence of M-CSF (20 ng/ml) only as a control (M0), pro-inflammatory medium supplemented with 20 ng/ml of lipopolysaccharide (LPS; Sigma Aldrich) and 20 ng/ml Interferon-*γ* (IFN*γ*; Peprotech) or anti-inflammatory medium supplemented with 40 ng/mL Interleukin 4 (IL-4; Peprotech) and 20ng/mL Interleukin 13 (IL-13; Peprotech). Polarized pro-inflammatory macrophages will be referred to as MΦ(LPS/IFN*γ*) and anti-inflammatory macrophages as MΦ(IL-4/IL-13).

### 2.3 qPCR

For RNA isolation, cells were washed three times with PBS and lysed with QIAGEN RLT lysis buffer (QIAGEN). Three wells per experimental condition were pooled for a total of 105,000 cells. RNA isolation was achieved following the protocol of the RNAeasy mini kit (QIAGEN) and RNA concentration was measured using a spectrophotometer (NanoDrop™ One/OneC microvolume UV-Vis spectrophotometer; Thermofisher). Complementary DNA (cDNA) was synthesized with a PCR thermal cycler (C1000 Touch, Bio-rad, Hercules, CA, USA). Macrophage polarization genes were analyzed with real-time PCR (CFX348 Touch Real-Time PCR detection system; Bio-Rad) and relative gene expression was quantified using the ΔCt method. The primer sequences are listed in Supplementary Table. **??**. GAPDH served as the internal housekeeping control gene.

### 2.5 Immunofluorescence Imaging

Cells were fixed with 3.7% paraformaldehyde (formalin 37%; 104,033.1000, Merck) for 20 mins at room temperature, permeabilized for 30 mins with 0.5% Triton-X-100 (108,643, Merck) in PBS, and blocked for 1 h with 4% goat serum and 1% Bovine Serum Albumin (BSA) in 0.05% PBS-Tween (PBS-T). After blocking, cells were incubated overnight at 4^*°*^C with two combinations of antibodies; rabbit anti-tubulin (1:200; ab18251, Abcam) and mouse anti-vimentin (1:500; ab20346, Abcam) or, rabbit anti-tubulin (1:200; ab18251, Abcam) and mouse anti-vinculin (1:300; V9131, Sigma-Aldrich). The next day, cells were washed with 0.05% PBS-T and incubated for 1 h at room temperature with Phalloidin-Atto 550 (1:200; 19083, Sigma-Aldrich) and the secondary antibodies; goat anti-rabbit IgG Alexa Fluor 488 (1:300; A11008, Molecular Probes) and goat anti-mouse IgM Alexa Fluor 647 (1:300; A21240, Molecular Probes). Cells were then washed twice with PBS-T, and nuclei were stained with 4’ 6-diamidino-2-phenylindole dihydrochloride (DAPI; 1:200, D9542, Sigma-Aldrich), mounted in Mowiol on glass slides and stored in the dark at 4^*°*^C until imaging. Samples were imaged using the inverted widefield microscope Leica DMi8 with a 40x/0.60 airy objective.

### 2.5 Morphometrics and Clustering

The morphological analysis was performed on the immunofluorescence images obtained as in section 2.4 for a total amount of 2232 cells (n= 757 M0, n=732 MΦ(LPS/IFN*γ*) and n=743 MΦ(IL-4/IL-13)). The cells were segmented using CellPose ^16^, previously trained on multiple macrophage IF images. Cell area, perimeter, circularity, roundness, and aspect/ratio (Table 1) were calculated using a custom Python routine. To further explore macrophage morphologies, a second custom Python pipeline has been developed in order to sort them into morphological clusters. The clustering algorithm used is the K-means method, one of the most widely used methods for cell clustering ^17,18^.

**Table 1:** Morphometric parameters and their formulas. Number of pixels (*N*_*p*_), Number of vertical/horizontal pixel sides (*N*_*h,v*_), Number of diagonal pixels (*N*_*d*_)

| Morphometric | Formula |
| --- | --- |
| Area ($A$) | $N_p$ |
| Perimeter ($P$) | $\approx N_{h,v} + \sqrt{2} N_d$ |
| Circularity ($C$) | $\frac{4\pi A}{P^2}$ |
| Roundness ($R$) | $\frac{4A}{\pi(\text{MajorAxis})^2}$ |
| Aspect Ratio ($A_r$) | $\frac{\text{MajorAxis}}{\text{MinorAxis}}$ |

### 2.6 Atomic Force Microscopy measurements and cluster assignment

Immediately before AFM measurements, the culture medium was removed and replaced with fresh RPMI 1640 medium without cytokines. Mechanical measurements of macrophages were performed using an atomic force microscope (Nanowizard 4; Bruker, USA) mounted on an inverted optical microscope (Olympus IX73), enabling both force spectroscopy and microrheology. This characterization was performed on ≈10 cells per sample, selecting morphologies from the most representative clusters. Samples were maintained at physiological temperature using a Petri dish heater (Bruker, USA), preheated to 37°C. This temperature was kept constant throughout all measurements. An MLCT-SPH1*µ*M-B rectangular cantilever probe with a 1 µm radius hemispherical tip (Bruker, MLCT-SPH1µM, USA) and a nominal spring constant of 0.02 N/m was used. The cantilever’s sensitivity and actual spring constant were calibrated in the cell culture medium at 37°C using the thermal noise method, as described by Hutter and Bechhoefer ^19^.

#### 2.6.1 Force Spectroscopy

Force spectroscopy was carried out using a setpoint force of 200 pN, a z-velocity of 2.0 *µ*m/s, and a z-range of 6.0 *µ*m, resulting in an average indentation depth of 200–500 nm. To ensure the acquisition of force spectroscopy curves over the perinuclear region, a 20 × 20 *µ*m area was first scanned, and an image was acquired. Then, a 10 *µ*m × 10 *µ*m force spectroscopy map was acquired in the nuclear region, recording force spectroscopy at 9 positions arranged in a 3 × 3 grid. Data analysis was performed using the JPK-SPM postprocessing software (Bruker, USA). The elastic modulus values in each scanned point were obtained by fitting the first 200 nm (indentation depth) of the approach segment of the force–distance curves to the Hertz model, considering a spherical indenter with the nominal tip radius value and a Poisson ratio of 0.5. The elastic modulus values for each cell were calculated as the average elastic modulus value in the region above the nucleus. In this region, the cell height was always larger than 6 *µ*m. Therefore, the Hertz condition, namely an indentation depth lower than 10% of the thickness (height) of the sample (cell), was satisfied, and no corrections for the bottom effect needed to be applied ^20^.

### 2.6.2 Microrheology

For microrheological analysis, the same 10 *µ*m × 10 *µ*m area within the perinuclear region was selected for each cell. The same setpoint force and z-velocity as used in force spectroscopy were applied, but the z-range was extended to 8 *µ*m. After the initial approach and a 1-second pause, the probe was oscillated with an amplitude of 25 nm for 20 cycles. Measurements were performed over the frequency range 1.6–250 Hz, using five log-scaled frequency points per decade. For each cell, four measurements were taken at different locations within the selected area. Microrheology data were processed using the JPK-SPM postprocessing software (Bruker, USA), following the method described by Alcaraz et al ^21^ which calculates the complex elastic modulus E based on the Fourier transform of the force and indentation signals. A Poisson’s ratio (*v*) of 0.5^22^, and a spherical indenter with the nominal tip radius value (1 *µ*m) was assumed. The frequency-dependent complex modulus data were fitted using a double-power-law (DPL) model ^23^ in Origin (OriginPro, Version2020).

### 2.6.3 Statistical Analysis

All datasets were tested for normality using the Shapiro-Wilk’s test. For the gene expression analysis, normality assessment and statistical tests were performed on the ΔCT values using all individual data points pooled across donors, without accounting for donor-specific effects. A One-way ANOVA with Tukey’s multiple comparisons test was performed on IL-6, and the non-parametric Kruskall-Wallis test with Dunn’s multiple comparisons test was performed on all the other genes (*p¡0.05; **p¡0.01; ***p¡0.001; ****p¡0.0001). For the morphological characterisation the data are shown as a superplot following the method of Lord at al. ^24^ in which data from all cells is superimposed with the mean SEM of each donor; n=3. The statistical analysis was performed on these means (i.e. n=3) using non-parametric Kruskall-Wallis test with Dunn’s multiple comparisons test. For the mechanical data, the non-parametric Kruskall-Wallis test with Dunn’s multiple comparisons test was performed on elastic modulus (E) and high frequency data (*β*) and a one way ANOVA with Tukey’s multiple comparisons test on the low frequency data (*α*) (*p¡0.05; **p¡0.01; ***p¡0.001; ****p¡0.0001). All statistical anyalyses and graphs were performed using GraphPad Prism version 11.0.0 for Windows, GraphPad Software, Boston, Massachusetts USA, www.graphpad.com.

## 3 Results

### 3.1 Gene expression analysis confirms macrophage polarization

The successful polarization of macrophages into pro-inflammatory and anti-inflammatory phenotypes was first confirmed by assessing the gene expression of *IL6, TNFα*, and *CCR7* as markers for MΦ(LPS/IFN*γ*), and *CD206* as a marker for MΦ(IL-4/IL-13) (Fig 1). qRT-PCR results showed a higher expression of *IL6, TNFα*, and *CCR7* for MΦ(LPS/IFN*γ*) compared to unstimulated M0 macrophages and MΦ(IL-4/IL-13) for all three donors. TGF*β* expression was similar between the three macrophages states. This could be explained by the fact that TGF*β* is an important regulatory mediator in the immune response, and while TGF*β* is often associated with alternatively activated macrophages, regulatory genes are often co-expressed in both polarizations as part of a feedback loop ^25^ . Finally, *CD206* expression was significantly higher in MΦ(IL-4/IL-13) compared to M0 macropahges and MΦ(LPS/IFN*γ*). From these results, we can conclude that macrophages were successfully polarized into MΦ(LPS/IFN*γ*) and MΦ(IL-4/IL-13).

**Figure 1:**
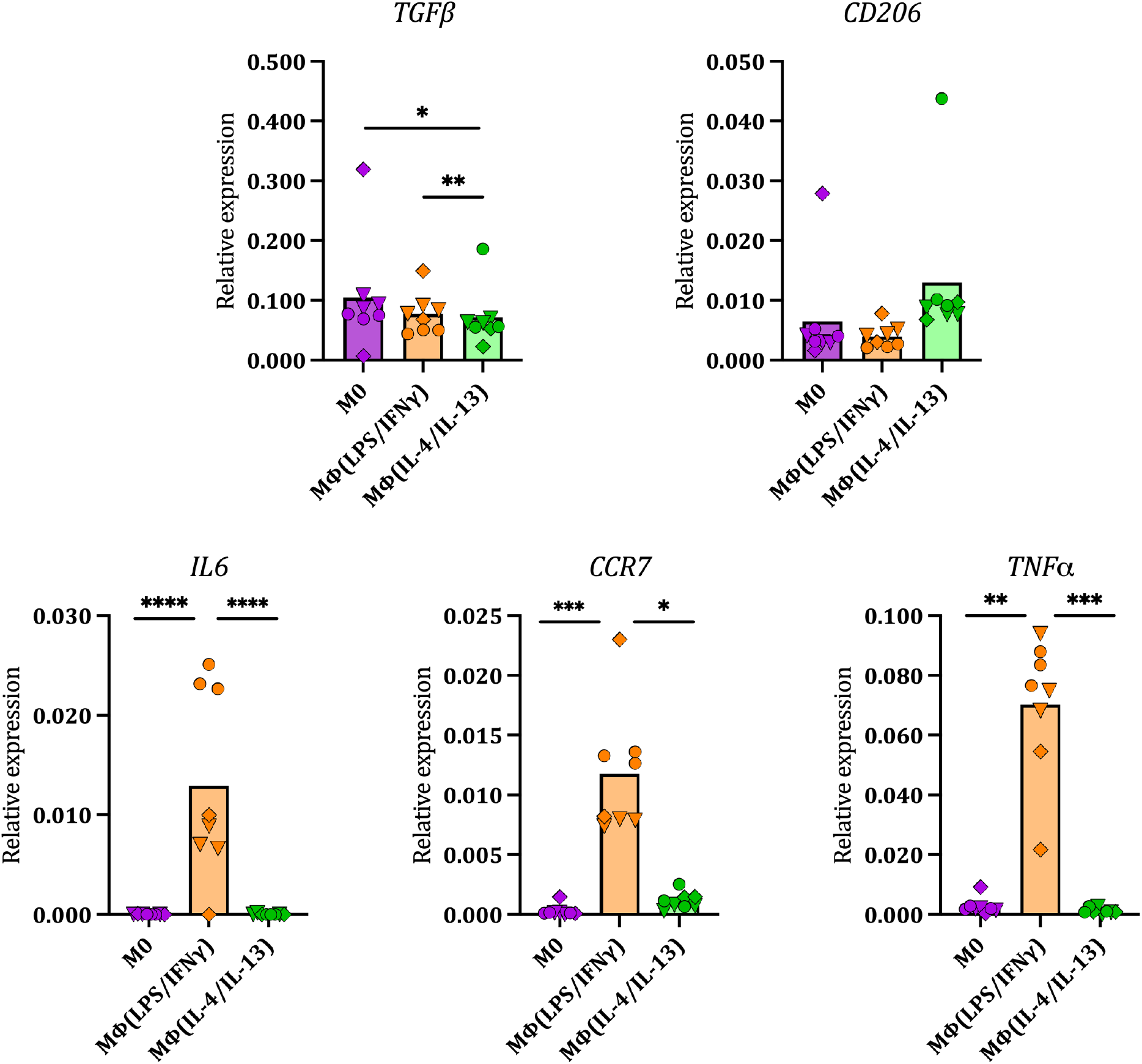
Relative expression of macrophage anti-inflammatory gene (CD206) and pro-inflammatory genes (IL6, TNF, and CCR7) in M0, MΦ(LPS/IFN*γ*) and MΦ(IL-4/IL-13) for 3 different donors (n=3). The polarization states are represented in purple for M0, orange for MΦ(LPS/IFN*γ*) and green for MΦ(IL-4/IL-13). Symbols denote individual donors; ▽ Donor 1, ◊ Donor 2, ◯ Donor 3, (n = 3) . For each condition, three points per donor are plotted, representing technical replicates from the well plate. Each point is the mean of two technical qPCR replicates. Statistical analysis was performed on the ΔCT values, on all individual points pooled across donors (*p¡0.05; **p¡0.01; ***p¡0.001; ****p¡0.0001)

### 3.2 Macrophages are morphologically heterogenous

A morphological analysis of macrophages under different polarization conditions was performed with two objectives. First, we wanted to assess the effect of polarization on macrophage morphology, which has been highlighted in previous studies ^5,26–29^. In addition, because macrophage morphology is highly heterogeneous and AFM is an intrinsically low-throughput technique, we sought to identify the most representative morphologies for each biological condition to guide AFM mechanical testing. For this purpose, immunofluorescence images of polarized and unpolarized macrophages from three donors were analyzed using different morphometric parameters (Fig 2). The results show no statistically significant differences in morphology between M0, MΦ(LPS/IFN*γ*) and MΦ(IL-4/IL-13) for all parameters. There was a tendency for an increase in eccentricity in MΦ(LPS/IFN*γ*) compared to M0 and MΦ(IL-4/IL-13), although this was not consistent across all three donors. Moreover, the spread of the data (Fig 2a) reflects a high heterogeneity in macrophage morphology within each population, which is also observed in the immunofluorescence images of the cells and their representative masks in Fig 2b. Therefore, in order to better understand the results, the acquired morphological data were further analyzed to generate morphological clusters. The clustering algorithm resulted in three different morphological groups (Fig 3a & Fig 3b) that we classified as; **(0)** round and small, **(1)** round and big, **(2)** elongated (Fig 5).The abundance of the three morphological groups showed a similar distribution across all three polarization states, with cluster **0** being the predominant cluster (≈71%-63%), followed by cluster **1** (≈31%-26%), and finally cluster **2**(≈5%-1.85%) (3c). Moreover, round cells (clusters **0** and **1**) represent on average 98%-95% of the total population in each polarized state. These results suggest that there is no clear relationship between morphology and macrophage polarization and that round cells constitute the majority of the population. Based on these results, we focused the AFM-based mechanical characterization only on clusters 0 and 1 (Supplementary Figure **??**).

**Figure 2:**
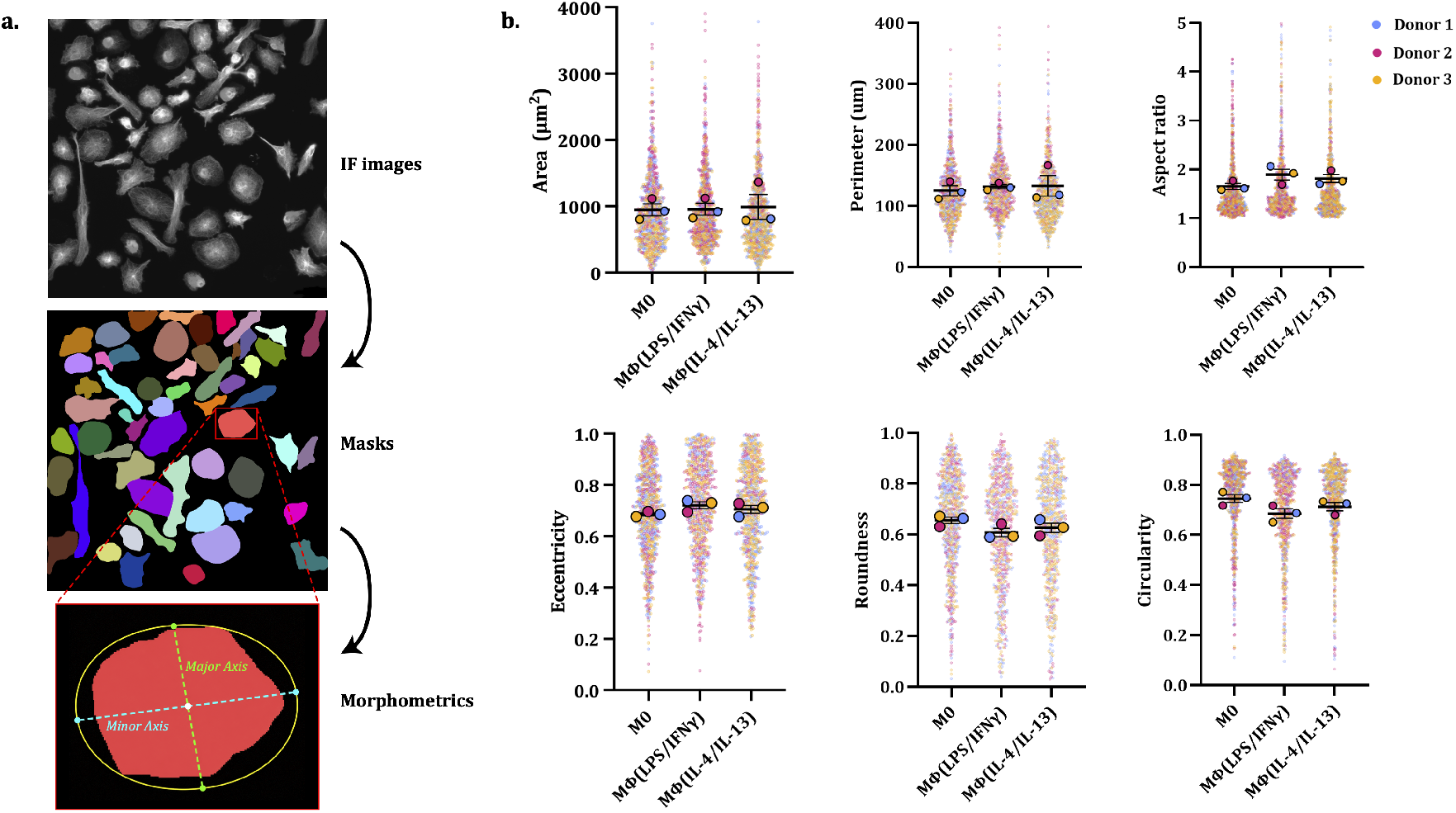
a. Image analysis pipeline to extract single cell morphometrics, with MΦ(LPS/IFN*γ*) displayed as an example b. Superplots showing the morphological characterization of M0, MΦ(LPS/IFN*γ*), and MΦ(IL-4/IL-13)) macrophages from 3 donors. Small dots represent individual cells, while larger dots represent the mean of each donor (n = 3 donors). Statistical analyses were performed on donor-level means using a Kruskal–Wallis test.

**Figure 3:**
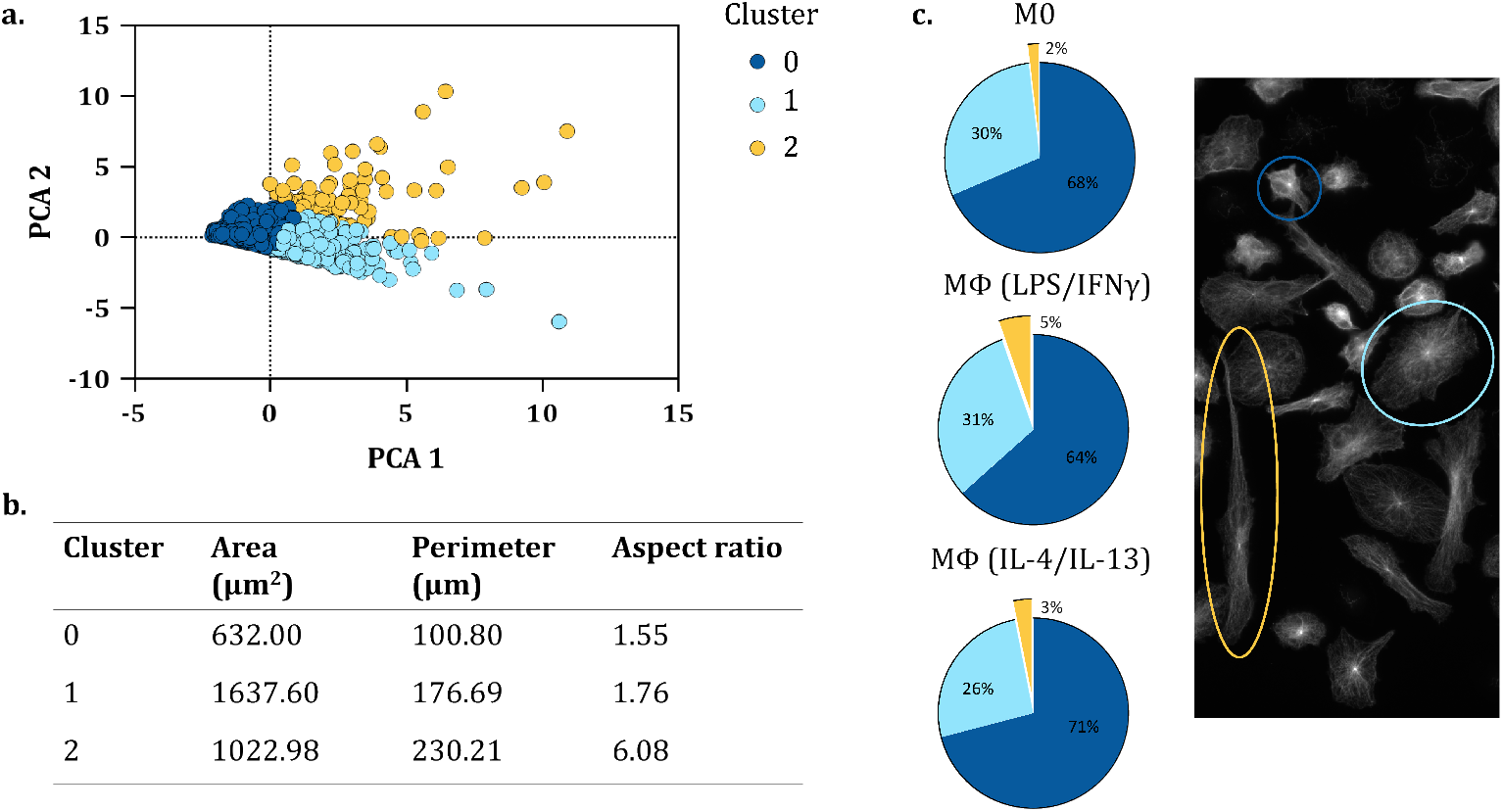
a. Principle Component Analysis (PCA) of the area, perimeter and aspect ratio of the analyzed cells b. Table summarizing the average of the three morphological parameters within each cluster c. Pie charts showing the distribution of the clusters for M0, MMΦ(LPS/IFN*γ*) and MΦ(IL-4/IL-13) and IF image showing an example of a cell belonging to cluster 0 (dark blue), cluster 1 (light blue) and cluster 2 (yellow)

### 3.3 Polarization alters macrophage mechanical properties

We performed AFM-based cell mechanical measurements on morphologically selected cells (10 cells with a non-elongated morphology per condition). The mechanical characterizations were performed on M0 as a control, and on MΦ(LPS/IFN*γ*) and MΦ(IL-4/IL-13) macrophages, representing pro-inflammatory and anti-inflammatory polarization, respectively, with an indentation depth in the range 200 - 500 nm. Therefore, the measured mechanical parameters are representative of the cortical region and reflect contributions from the mechanical properties of the membrane and cytoskeleton. The average elastic modulus of pro-inflammatory macrophages was significantly lower than the elastic modulus of anti-inflammatory macrophages as measured by AFM force spectroscopy (458.8 ± 179.9 Pa and 716.0 ± 345.6 Pa, respectively). The average value of the control (unstimulated) M0 macrophages was similar to that of anti-inflammatory macrophages (754.6 ± 367.2 Pa), but due to the higher spread of the data, it did not show any significant difference when compared to pro-inflammatory macrophages (Fig 4a).

**Figure 4:**
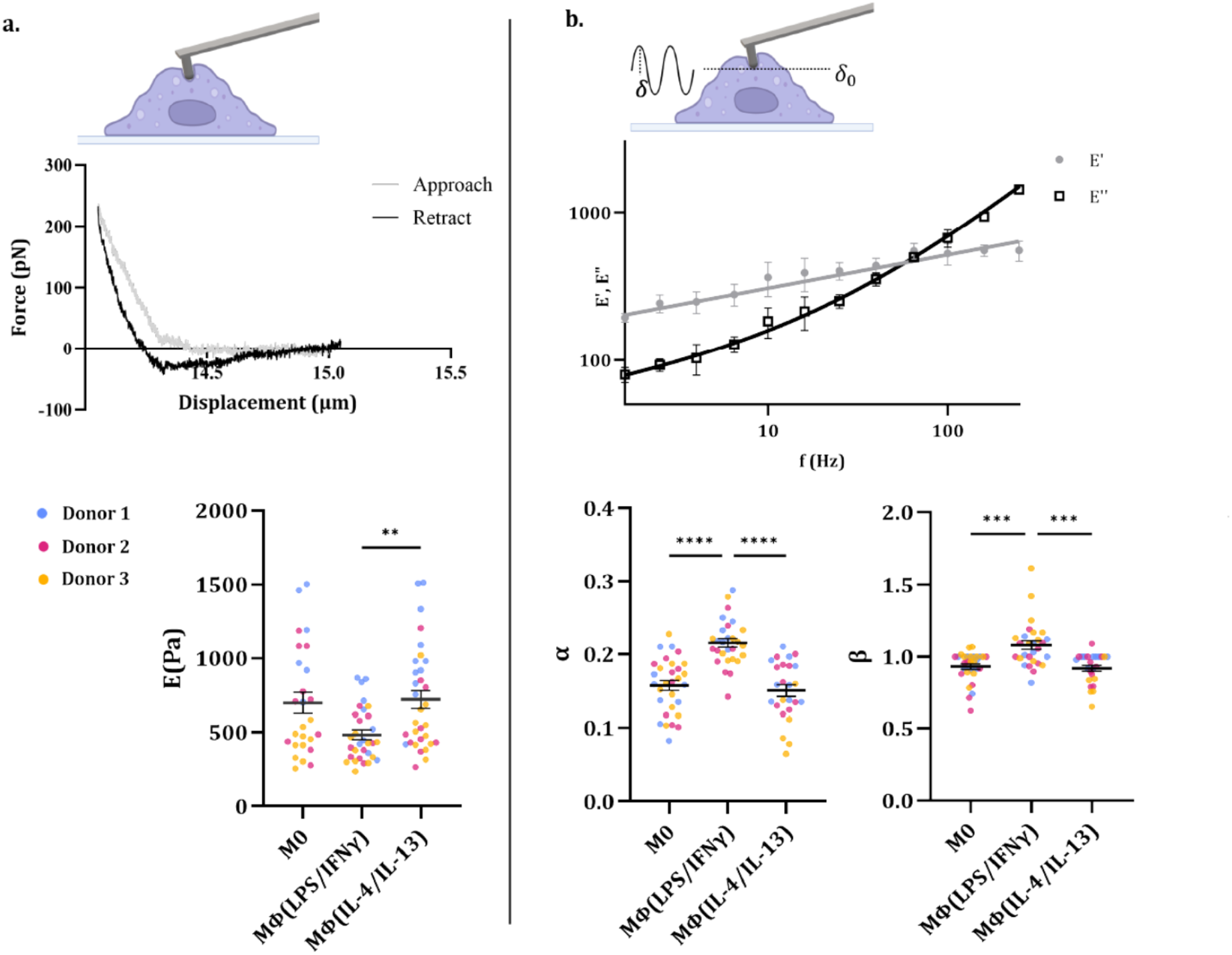
a. Force spectroscopy measurements: schematics of the experimental procedure (top) (Biorender.com), example of force-displacement curve (center), and Elastic Modulus “E” of M0, MΦ (LPS/IFN*γ*) and MΦ (IL-4/IL-13) macrophages (bottom). b. Microrheology measurements: schematics of the experimental procedure (top) (Biorender.com), example of frequency sweep curve (center), and values of the fitted parameters *α* and *β* of M0, MΦ (LPS/IFN*γ*) and MΦ (IL-4/IL-13) macrophages (bottom). A Kruskal-Wallis test was perfromed on the elastic modulus and the high frequency data (*β*) and a one way ANOVA with Tukey’s multiple comparisons test on the low frequency data (*α*) (**p¡0.01; ***p¡0.001; ****p¡0.0001))

AFM-based microrheology measurements revealed that, under the measured conditions, macrophage behavior in the analyzed frequency range (1-250 Hz) is well described by a double power law (DPL) model (Fig 4b), showing distinct regimes at low and high frequencies (Fig 4b), as with other types of mammalian cells previously investigated with similar methods ^23^. The DPL corresponds to the fractional Kelvin-Voigt framework, which differs from the classical Kelvin-Voigt model by using springpot elements rather than standard springs and dashpots. Springpots, depending on the value of the fractional exponent, can behave with characteristics between those of an ideal spring (power exponent = 0) and a dashpot (power exponent = 1). Fitting the microrheology data to the DPL model yielded the low- and high-frequency power-law exponents (*α* and *β*, respectively), which describe the cell’s viscoelastic response in the low- and high-frequency regimes, respectively. The model is expressed as:

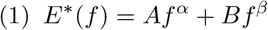

where A and B are constant values of amplitude in the low- and high frequency regimes, respectively. The corresponding real component (storage modulus E’) and the imaginary component (loss modulus E”) are defined, respectively, as:

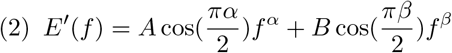

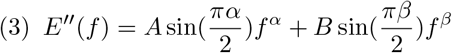

Fitting the microrheology data to the DPL model showed that all analyzed cells exhibited two response regimes in the analysed frequency range, characterized by a dominant elastic solid-like behavior (E’¿E”) at low frequency (long timescale) and a dominant dissipative, fluid-like behaviour (E”¿E’) at high frequency (short timescale), described by low- and high-frequency power-law exponents (*α* and *β*, respectively). This behavior is compatible with the biphasic and active nature of live cells, where cortical cytoskeleton tension and contractility dominates the long time-scale and the cytoplasm flow and friction dominate the short timescale response ^30,31^.

In our experimental conditions, cells exhibited low-frequency exponents in the range 1.6-120 Hz and high-frequency exponents in the range 120-400 Hz, close to 1, confirming dominant elastic behavior at low frequencies and a more dissipative response at high frequencies. The frequency switch between these two modes is usually expressed as crossing time *τ* (Supplementary Figure **??**). The observed high values of power law exponents have previously been associated with actin depolymerization and transient crosslinking ^23,32^, suggesting a low degree of actin polymerization across all analyzed cell types. Physically meaningless values of *β* above 1 have been observed elsewhere and ascribed to the contribution of additional components in the cell’s viscoelastic response, which are not well described by the double power law model ^33^. Anti-inflammatory macrophages exhibited significantly lower values of both low- and high-frequency exponents (*α* = 0.15± 0.04 and *β* = 0.91 ± 0.11) compared to pro-inflammatory macrophages (*α* = 0.22 ± 0.03 and *β* = 1.07 ± 0.16), which indicates a more elastic and less dissipative response for M2-like cells in the entire tested frequency range. Low values of power-law exponents have previously been attributed to high cytoskeletal tension, pre-stress, and contractility ^23,32,34^. Therefore, the results suggest a higher tension and a more contractile state for anti-inflammatory macrophages compared to pro-inflammatory ones.

### 3.4 Polarized macrophages exhibit distinct podosomes distributions

To determine whether the differences in the measured mechanical properties of MΦ (LPS/IFN*γ*) and MΦ (IL-4 / IL-13) are associated with cytoskeletal changes, we stained polarized macrophages for the main components of the cytoskeletal network: tubulin, F-actin, and vimentin. Figure 5a and b show representative immunofluorescence images of MΦ (LPS/IFN*γ*) and MΦ (IL-4 / IL-13) for each cytoskeletal protein. No clear differences can be observed in microtubuli, and vimentin structures and expression between MΦ (LPS/IFN*γ*) and MΦ (IL-4/IL-13) (Fig 5 a,b). However, while the actin staining did not show differences in fiber distribution between polarization states, we could observe a difference in spatial distribution of actin-rich dots, that we identified as podosomes (Fig 5 c,d). Podosomes are dynamic adhesion structures found in myeloid cells, with an important role in macrophage adhesion and migration with additional roles in mechanosensing, matrix degradation, cell-cell fusion and phagocytosis. Their structure is characterized by an actin rich core surrounded by a ring of integrins, vinculin, and other adhesion adaptor proteins ^35^ . Therefore, to further investigate the differences in podosome distribution between polarized macrophages, we imaged cells for both vinculin and actin. In anti-inflammatory macrophages, podosomes were organized in well defined clusters, and often localized at the leading and/or rear edge (5 c). In contrast, pro-inflammatory macrophages exhibited a more dispersed and sparse podosome distribution throughout the cell (5 d). Additional images of podosomes can be found in Supplementary Figure **??**.

**Figure 5:**
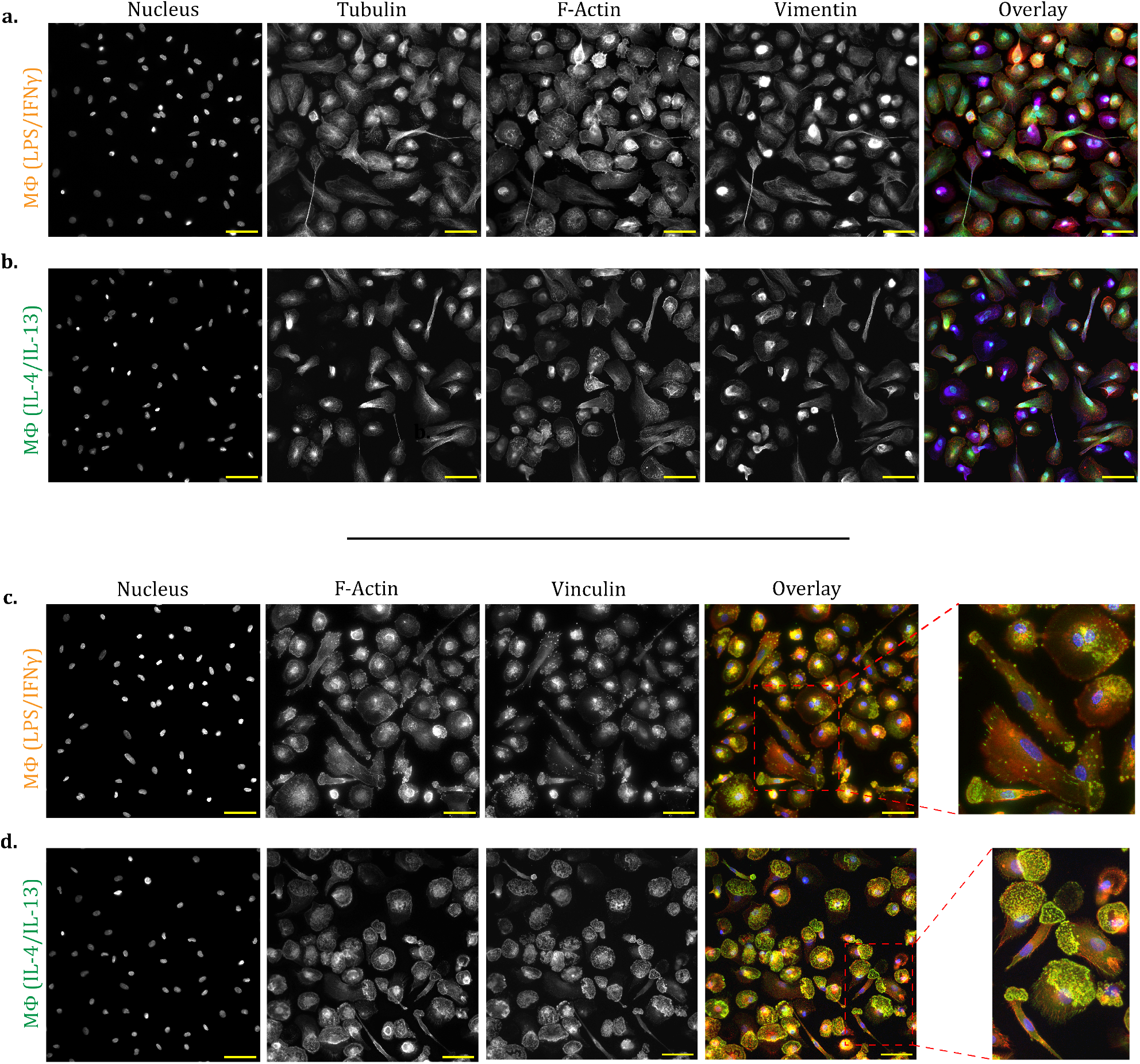
Representative images of tubulin, F-actin and vimentin organization in (a) MΦ (LPS/IFN*γ*) and (b) MΦ (IL-4/IL-13) macrophages. The overlay shows the nucleus/DAPI in cyan, tubulin in green, F-actin in red, vimentin in dark blue. Representative images of F-actin and vinculin stainings to visualize podosome organiztion in (c) MΦ (LPS/IFN*γ*) and (d) MΦ (IL-4/IL-13) polarized macrophages. The overlay shows the nucleus/DAPI in dark blue, F-actin in red and vinculin in green. Scale bars: 50 µm

## 4 Discussion

Macrophages are phenotypically plastic cells that dynamically adapt their morphology and function in response to changes in their microenvironment, which makes them challenging cells to study. In this work, we first investigated the possible link between function and morphology with the double aim of: analyzing the possibility of using morphometric parameters as markers for macrophage phenotype; identifying the most representative cell morphologies for each analyzed cell population to focus the (intrinsically low-throughput) AFM mechanical analysis on those cell morphologies. Under our experimental conditions, we could not observe a direct relationship between morphology and biochemically induced phenotype; the different populations exhibited large morphological heterogeneity, as reflected in the overlap in the quantified morphometric data. This is consistent with previous studies, in which large standard deviations and overlap of morphometric parameters among different population groups were also observed ^26,36^. As an example, Malheiro et al. reported an increase in cell spread for IL-4-stimulated THP-1-derived macrophages compared to the LPS/IFN*γ*-stimulated ones, but no significant differences in circularity and aspect ratio ^36^. Similarly, Selig et al. observed high morphological heterogeneity in different hPBMC-derived macrophage phenotypes and succeeded in defining an AI-based prediction model only by introducing a synthetic dataset generated from single-cell data ^26^. Altogether, these results indicate that no univocal relationship exists between biochemically induced hPBMC-derived macrophage phenotype and morphology.

AFM-based mechanical characterization at the single cell level, especially the frequency sweep for microrheological measurements is a slow procedure. Each of our sample was measured under controlled temperature conditions, within a 2 hour time frame, to avoid cell damage and death. Due to this time constraint, the sample size was reduced to around 10 cells per donor per condition. The small sample size does not allow us to fully capture the sample’s morphological heterogeneity. Therefore, we restricted the analysis to cells that share a similar morphology. Our analysis allowed us to identify the most abundant cell morphometric clusters for each population, enabling selection of the most representative cell morphology for AFM mechanical analysis. Specifically, for all three analyzed macrophage populations, about 95% of cells exhibited on average an aspect ratio between 1.55 and 1.76; and elongated cells (average aspect ratio of 6.08) accounted for less than 5% of the total populations and were therefore excluded from the mechanical analysis.

Interestingly, our AFM mechanical characterization of primary human monocyte-derived macrophages showed that differently polarized macrophages exhibit distinct mechanical properties, with the pro-inflammatory phenotype exhibiting a lower elastic modulus (458.8 ± 179.9 Pa vs. 716 ± 345.6 Pa) and higher viscosity as reflected by higher values of low- and high-frequency components (*α* = 0.22 ± 0.03 vs. *α* = 0.15 ± 0.04 and *β* = 1.07 ± 0.16 vs *β* = 0.91 ± 0.03) compared to the anti-inflammatory phenotype. This resulted independently of cell morphology. Unfortunately, direct comparisons with other studies on the mechanics of primary macrophages are impossible due to substantial experimental differences. As an example, Evers et al., using optical tweezers, also observed a lower elastic modulus for hPBMCs-derived pro-inflammatory macrophages (≈650 Pa vs. ≈750 Pa) and a higher power-law exponent (0.48 ± 0.07 vs. 0.29 ± 0.04) compared to anti-inflammatory macrophages, consistent with our results. However, it is worth noting that both biological and mechanical conditions were consistently different from those in our study, which might explain the (slight) quantitative discrepancies. Particularly, the main experimental differences include: i) macrophage phenotypes were differently stimulated (GM-CSF was used to induce a pro-inflammatory phenotype, while M-CSF was used to stimulate the anti-inflammatory phenotype); ii) after culture, the cells were analyzed in a (re-)suspended state; iii) the level of strain and the velocity at which strain was applied were in a different range.

In another study, Suleimanov et al. investigated the differences in morphology and mechanics between four polarization states of hPBMCs-derived macrophages. Two M0 states; MΦ(M-CSF) and MΦ(GM-CSF), as well as an anti-inflammatory “M2” state (M-CSF, IL-4) and a pro-inflammatory “M1” state (GM-CSF, LPS + IFN*γ*) in the presence of phorbol-12-myristate-13-acetate (PMA), particulate opsonized zymosan (OZ) or E.Coli. For the mechanical characterization, they obtained mechanical maps of macrophages using AFM-based fast force volume mode, which involves acquiring force curves across the cell surface with high speed (specifically, 183 *µ*m/s) and fitting the FS curves with Hertz and Ting models. Although efficient at obtaining high-resolution information on the spatial variability of cell elastic modulus and viscosity within a single cell, the fast force volume technique yields cell elastic modulus values that are affected by indentation speed and, therefore, are significantly higher than those we obtained (at an indentation speed of 2 *µ*m/s). As for the viscoelastic measurements, Suleimanov et al. used a static loading of the cells, which is not comparable to our frequency-domain analysis ^12^.

Finally, Chen et al. proposed an AFM-based mechanical characterization of three phenotypes of monocytes-derived human macrophages (M0, MΦ(LPS/IFN*γ*) and MΦ(IL-10)). They employed force-volume mode, which allows for cell surface mapping at a lower indentation speed than the afore-mentioned fast-force volume mode. Although no statistically significant differences were observed, the cell elastic modulus was slightly higher in pro-inflammatory macrophages than in anti-inflammatory ones, apparently in contrast with our findings. However, the anti-inflammatory phenotype included in their study was polarized with Interleukin 10 (IL-10) making the results not comparable with ours.

Rheological studies of living cells provide information about the internal complexity of the cellular structure. Similar to other previously investigated cell types, primary human monocyte-derived macrophages exhibited a distinctive viscoelastic response to stimulation at low and high frequencies, which is well described by the DPL model and attributable to the cells’ active and biphasic nature. The cell cytoskeleton’s ability to remodel is strongly linked to the timescale or, in this case, the frequency of the stimulation. A long timescale allows for turnover of cytoskeletal proteins and binding/unbinding of crosslinkers, thereby allowing the cytoskeleton to rearrange and properly flow. At high frequency (short timescale), the ability of cytoskeletal crosslinkers to rearrange is limited, the cytoplasm becomes compressed, and, aside from an apparent stiffening of the cell, we observe an increase in dissipation of the stimulus throughout the cell structure ^30,31^. The differences in mechanics we observe between MΦ(LPS/IFN*γ*) and MΦ(IL-4/IL-13), therefore, likely reflect structural differences between these two phenotypes. Specifically, a more elastic-solid response (lower power-law exponent) at long time scales (low frequency), as we observed for anti-inflammatory macrophages compared to pro-inflammatory macrophages, has previously been attributed to higher cortical tension and cytoskeletal contractility ^23,32,34^., However, the structural composition of the cytoskeleton in human macrophage phenotypes is not well characterized in the literature. Therefore, we attempted to compare the cytoskeletal differences using IF staining. While no apparent differences were observed in tubulin, actin and vimentin fiber organization within the cell structure, vinculin and actin stainings showed distinct podosomes distribution in pro- and anti-inflammatory macrophages, consistent with the results of our mechanical analysis. Indeed, MΦ (IL-4/IL-13) were characterized by organized clusters of podosomes localized at the cell edge, suggesting an actin-rich, protrusion-based mesenchymal migration mode for these cells ^37,38^, typical of anti-inflammatory macrophages. On the contrary, MΦ (LPS/IFN*γ*) exhibited a more diffuse and homogeneous distribution of podosomes throughout the cell, consistent with their lower motility ^38^. Although for macrophages the link between mechanical properties and cell motility has not been directly investigated, for other cell types, e.g. neutrophils, it has been shown that migratory states are characterized by a high, not uniformly distributed membrane tension (specifically, higher at the front edge and lower at the rear) compared to more quiescent states ^39–41^ with a lower and uniformly distributed membrane tension. In addition, Labernardie et al. studied the mechanical properties of podosome structures in macrophages on patterned substrates using protrusion force microscopy. These organized structures were found to have a significantly higher elastic modulus than podosome-free areas. Furthermore, they measured oscillations in podosome stiffness over time, which they linked to actin turnover and myosin contractility. These results support the idea that cells with different podosome distributions exhibit distinct mechanical properties, with podosome-rich areas displaying a stiffer actin cytoskeleton than podosome-free areas ^42^. Furthermore, previous 2D traction force microscopy measurements by Hind et al. show that anti-inflammatory macrophages exert significantly higher traction forces on their substrate compared to pro-inflammatory macrophages, confirming increased cytoskeleton contractility. Indeed, these differences in force generation between pro- and anti-inflammatory macrophages are linked to myosin-driven contractility. Inhibition of ROCK, which mediates actomyosin contractility, significantly reduced force generation in unstimulated and anti-inflammatory macrophages; however, no effect was observed in pro-inflammatory macrophages. In contrast, inhibition of acto-myosin polymerization with Blebbistatin resulted in a significant reduction in traction forces across all polarization states ^43^. These findings suggest that anti-inflammatory macrophages rely on both ROCK-dependent and ROCK-independent pathways to generate contractile forces, whilst pro-inflammatory macrophages predominantly rely on a ROCK-independent pathway. One potential ROCK-independent mechanism involves myosin light chain kinase (MLCK), which can also promote myosin activation and contractility. Studies in fibroblasts and U2OS osteosarcoma cells suggest that ROCK- and MLCK-dependent myosin contractility are responsible for the organization and mechanical properties of stress fibers. These pathways differentially regulate the organization and mechanical properties of stress fibers. Specifically, ROCK-dependent pathways primarily regulate centrally located stress fibers, while MLCK activity controls peripheral stress fibers. Together, these observations suggest that distinct contractility pathways may underlie the polarization-dependent mechanical behavior of macrophages, and may also promote differences in actin cytoskeletal organization, with varying contributions of ROCK- and MLCK-mediated signaling potentially shaping the architecture and distribution of actin structures in each polarization state. ^44–46^. Specifically, the dependence of pro-inflammatory macrophages on ROCK-independent myosin contractility ^43^, along with the sparse podosome distribution, may lead to reduced actin polymerization in the perinuclear region and help explain the reduced elastic modulus observed in our cells.

### 4.1 Conclusion

In this study, we performed a micro-rheological analysis of different phenotypes of primary human monocyte-derived macrophages. This analysis showed that different phenotypes exhibited different elastic and viscoelastic signatures, which were found to be independent from cell morphology. Specifically, pro-inflammatory macrophages exhibited a lower elastic modulus and more dissipative behavior compared to anti-inflammatory macrophages, consistent with the predominant podosome distribution pattern observed in each phenotype, with clustered distributions in anti-inflammatory and sparse distributions in pro-inflammatory macrophages. Together, our results highlight cell mechanics as an integral component of macrophage phenotype, and provides a quantitative experimental framework to further unravel macrophage mechanobiology at the single-cell level. The 2D in vitro experimental setup used in this study provided a simple and well-controlled platform to perform fundamental measurements of macrophage activation and mechanics. Future studies would benefit from coupling the 2D cell mechanics approaches presented here with complementary investigations of macrophage mechanics in 3D environments. In particular, emerging techniques such as Brillouin microscopy and acoustic microscopy could provide additional insights into the mechanical properties of macrophages and their surrounding microenvironment in physiologically relevant 3D settings.

## Supplementary

Link to supplementary

## Author contributions

## Data availability

The datasets generated for this study can be found in the 4TU.ResearchData repository (doi: 10.4121/69d91f1f-7d98-431d-b867-212ebd3660e4).

## Acknowledgements

We thank Dr. Maja Rücker for her support in enabling the use of the AFM and the Multi-scale Laboratory of the Mechanics of Materials group at Eindhoven University of Technology. Microscopy was performed at the LCTE Microscopy Facility, TU/e. This research was financially supported by the European Research Council, ERC Starting Grant, Project 101042538 “MACxercise”. Views and opinions expressed are however those of the author(s) only and do not necessarily reflect those of the European Union or the European Research Council. Neither the European Union nor the granting authority can be held responsible for them. We gratefully acknowledge the Gravitation Program “Materials Driven Regeneration”, funded by the Netherlands Organization for Scientific Research (024.003.013).

