## Supplemental Information for "Polarization affects viscoelasticity and podosome distribution of primary human macrophages"

### Supplemental document

#### 1. SUPPLEMENTAL FIGURES

| | Age | Sex | Blood Type | Analysis |
|---|---|---|---|---|
| **Donor 1** | 74 | F | A+ | Morphology |
| **Donor 2** | 44 | M | A+ | Morphology |
| **Donor 3** | 59 | M | A+ | Morphology |
| **Donor A** | 67 | F | O+ | AFM |
| **Donor B** | 63 | M | A+ | AFM |
| **Donor C** | 65 | F | O+ | AFM |

**Fig. S1.** Information on the donors used in the experiments

| Gene name | Primer set | Code '5-'3 | Amplicon size (bp) | Position | Q-PCR |
|---|---|---|---|---|---|
| *IL-6* | Fw | GGTGAAGGTCGGAGTCAACG | 191 bp | 85-275 | E=100.2% |
| | Rev | GCTTCCCGTTCTCAGCCTT | | | |
| *B2M* | Fw | TAAGTGGGATCGAGACATGTAAG | 200 bp | 369-568 | E=100% |
| | Rev | GGAGACAGCACTCAAAGTAGAA | | | |
| *IL-6* | Fw | ACTCACCTCTTCAGAACGAATTG | 149 bp | 259-307 | E=97% |
| | Rev | GTCGAGGATGTACCGAATTTGT | | | |
| *CCR7* | Fw | AAGCCTGGTTCCTCCCTATC | 106 bp | 245-350 | E=101.7% |
| | Rev | ATGGTCTTGAGCCTCTTGAAATA | | | |
| *TNFα* | Fw | GAGGCCAAGCCCTGGTATG | 91 bp | 731-821 | E=101% |
| | Rev | CGGGCCGATTGATCTCAGC | | | |
| *CD206* | Fw | TGGGTTCCTCTCTGGTTTCC | 114 bp | 3920-4033 | E=102,7% |
| | Rev | CAACATTTCTGAACAATCCTATCCA | | | |

**Fig. S2.** qPCR primers

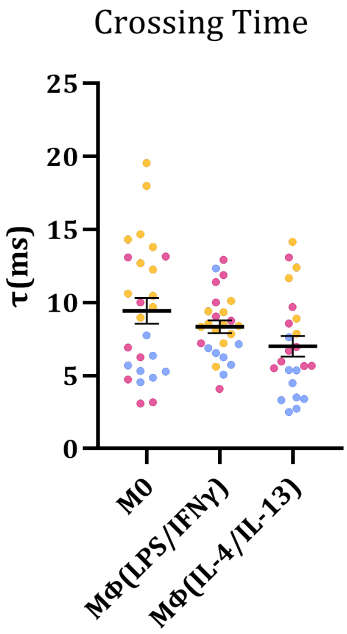


**Fig. S3.** Crossing time "Tau" ($\tau$), an additional parameter for the viscoelastic behavior of the cells for the three donors

Figure S3 $\tau$ is also an indicator of the viscoelastic behavior of the cell, being a descriptor of the relaxation time of the material; however, it has some intrinsic limitations. As it is obtained from the crossing point between the curves of storage and loss moduli over frequency, the value of $\tau$ depends on both the resting state of the cell (i.e., baseline) and its behavior over frequency. This means that this parameter is the most sensitive to the heterogeneity of the sample.

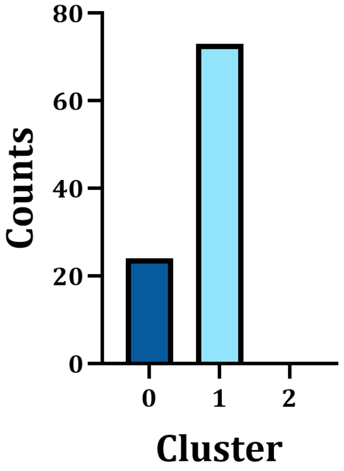


**Fig. S4.** Assignment of the cells measured during the mechanical characterization to the morphological clusters. The assignment is based on the calculation of the distance of each cell's morphological parameters with the cluster centroids.

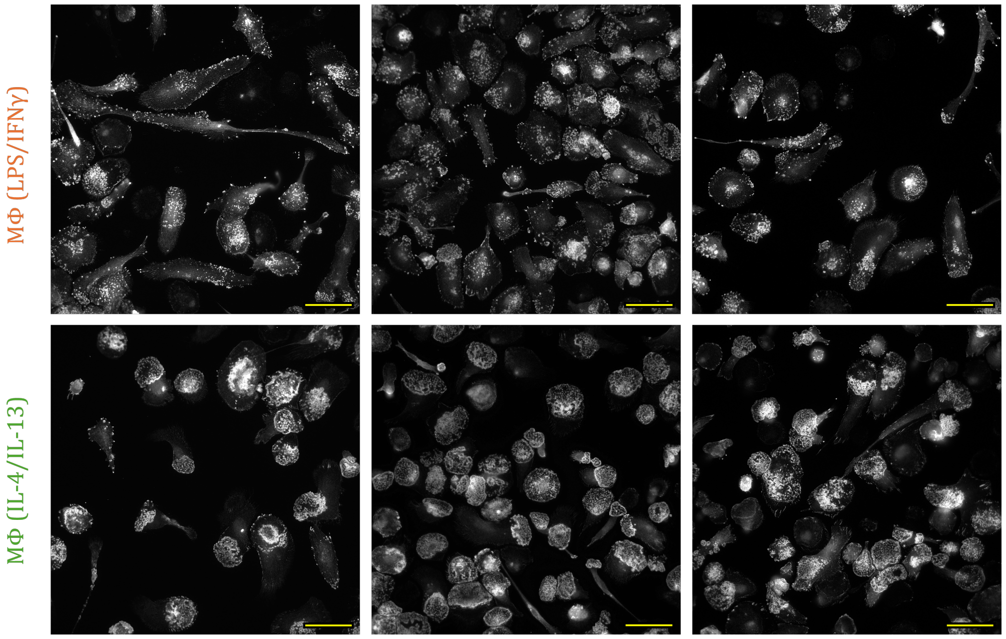


**Fig. S5.** Immunofluorescence images of MΦ(LPS/IFN$\gamma$) and MΦ(IL-4/IL-13) podosomes, stained with vinculin. Scale bar = 50$\mu$m
